# Rapid and Visible RPA-Cas12a fluorescence Assay for Accurate Detection of Zoonotic Dermatophytes

**DOI:** 10.1101/2021.06.04.446987

**Authors:** Liyang Wang, Jinyu Fu, Guang Cai, Di Zhang, Shuobo Shi, Yueping Zhang

## Abstract

**Background:** Dermatophytosis is an infectious disease of global significance caused by several fungal species, which affects the hair, nails, or superficial layers of the skin. The most common zoonotic dermatophytes are *Microsporum canis, Nannizzia gypsea* and *Trichophyton mentagrophytes*. Wood’s lamp examination, microscopic identification and fungal culture are the main conventional diagnostic methods used in clinics. Less common methods are dermatophyte PCR and biopsy/histopathology. However, these methods also have limitations for providing both accuracy and timely on-site detection. The recent development of CRISPR-based diagnostic platform provides the possibility of a rapid, accurate, and portable diagnostic tool, which has huge potential for clinical applications.

**Objectives:** The purpose of this study is to establish a molecular method for rapid and accurate diagnosis of clinical dermatophytes, which can accelerate clinical diagnostic testing and help timely treatment.

**Methods:** In this paper, we design a Cas12a-based assay combined with recombinase polymerase amplification (RPA) to differentiate three main zoonotic dermatophytes. The limit of detection (LOD) is determined by using standard strains. A total of 25 clinical samples (hair and scurf) are identified to evaluate the sensitivity and specificity of this assay.

**Results:** The RPA-Cas12a method showed high sensitivity and specificity (100% and 100%, respectively). The results could be observed directly by naked-eyes, and all tested samples were consistent with fungal culture and sequencing results.

**Conclusions:** Compared with other methods, the RPA-Cas12a-fluorescence assay requires less time (30 minutes) and less complicated equipment, and visible changes can be clearly observed, which is suitable for on-site clinical diagnosis.

## 1 INTRODUCTION

Dermatophytosis is an infectious disease of global significance caused by several fungal species, which affects the hair, nails, and/or superficial layers of the skin.^1^ The pathogens of dermatophytosis are called dermatophytes, which can be zoophilic, geophilic or anthropophilic fungal organisms in small animals. The most common zoonotic dermatophytes are *Microsporum canis, Nannizzia gypsea* (also called *Microsporum gypsea*) and *Trichophyton mentagrophytes*.^2^ In cats, 98% of infections are caused by *M. canis*^3^ Dermatophytosis is acquired by contact with infectious substances (i. e. spores, hyphae) through direct animal-to-animal contact or exposure to the soil.^2^ Clinical appearances are extremely variable, including hair loss, papules, erythema, scaling, crusting, and hyperpigmentation. Pruritus can be present or absent. There is no gold standard diagnostic method for dermatophytosis, and diagnostic procedures should always be guided by the clinical history of patients and clinician’s suspicion of diseases.^4^ The most commonly used conventional diagnostic methods for dermatophytosis are Wood’s lamp examination, microscopic identification and fungal culture. Less common methods are dermatophyte PCR and biopsy/histopathology. Microscopic cytology findings can help clinicians start initial treatment before culture results obtained. Molecular tools have been increasingly developed for dermatophytes detection (i. e. conventional PCR,^5,6^ real-time PCR,^7^ nested PCR,^8^ multiplex PCR,^9^ PCR enzyme-linked immunosorbent assay^10^ and PCR-restricted fragment length polymorphism^11–14^). PCR has shown the potential to detect the pathogens in mixed infections or environmental fungal contamination within about 2-3 hours,^15^ yet it is more reliant on available laboratory conditions and facilities.^16^ MALDI-TOF (matrix-assisted laser desorption/ionization time-of-flight) mass spectrometry (MS) has also been reported in dermatophytes identification,^17^ which is limited by the database availability and facility price. The identification of dermatophytes can help veterinarians to implement anti-fungal therapy on time and go a step further in determining the source of infection. What’s more, identifying specific pathogens can help veterinarians to educate the owners for effective prevention according to different transmission routes of different pathogens, which can effectively prevent the disease from recurring.

CRISPR-Cas12a protein has been especially noted in molecular diagnostics in recent years. The single multi-domain protein Cas12a (also called Cpf1), which is classified as a Class 2 nuclease, has the activity of inducing RNA-guided dsDNA cleavage.^18,19^ Like CRISPR-Cas9, the Cas12a protein has been harnessed for genome editing. However, recent studies also find that Cas12a protein has a target-activated, non-specific single-stranded deoxyribonuclease (ssDNase) cleavage activity. Briefly, after crRNA specifically hybridized with target dsDNA, Cas12a is activated to have indiscriminate cleavage activity to degrade ssDNA or RNA. Meanwhile, with the help of a fluorophore quencher (FQ)-labeled reporter, the results are able to be observed by naked-eyes with blue light. When performing this technique, isothermal pre-amplification steps are usually combined for the target DNA enrichment, which is less costly than PCR-based equipment requirements. Recently, based on the process of achieving attomolar sensitivity in the target DNA detection, Chen *et al*. created the DNA endonuclease-targeted CRISPR trans reporter (DETECTR).^20^ On the basis of this discovery, they reported and applied Cas12a protein by combining recombinase polymerase amplification (RPA) to develop a rapid and accurate test for the detection and classification of human papillomavirus (HPV) in clinical specimens.^20^ Based on the ssDNase property of the Cas12a protein, it expands upon the diagnostic applications range of infectious and noninfectious diseases in various situations, such as the detection of virus (i. e. pandemic COVID-19^21^, SARS-CoV-2^22^, and African swine fever virus^23^) and bacteria^24^ (i. e. *Staphylococcus aureus*^25^ *Listeria monocytogenes*^26^ *Mycobacterium tuberculosis*^27^, and Methicillin-resistant *Staphylococcus aureus* (MRSA)^28^), it can also be used for pathogen detection in agricultural^29^ and aquatic community.^30^ CRISPR technology is expected to become a rapid, accurate and portable diagnostic tool in the future.^31^

It has been noted that an accurate, rapid, and user-friendly assay is desired in small animal clinics to solve the shortcomings of conventional dermatophytes diagnostic methods. Meanwhile, CRISPR technology has shown the potential for rapid and accurate detection of nucleic acids. In this study, we firstly establish a method for detecting dermatophytes using the CRISPR-Cas12a based detection combined with the RPA reaction. Compared with other methods, the Cas12a-based assay requires less time and less complicated equipment with high sensitivity and specificity (Fig. 1). After incubation at room temperature, visible changes can be clearly observed. In conclusion, this research provides a new method for the detection and identification of dermatophytes and will improve the efficiency of on-site clinical diagnosis.

**Fig. 1:**
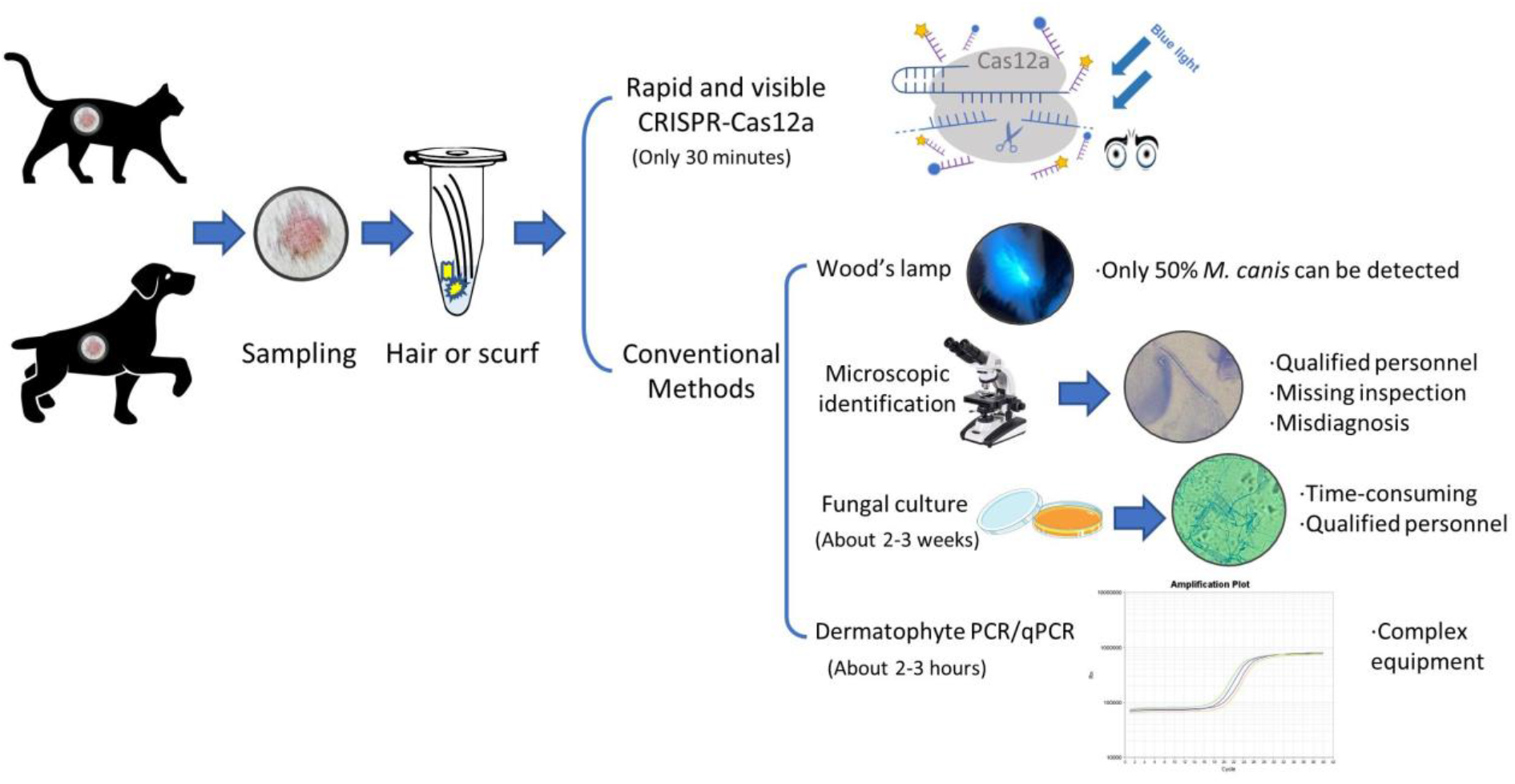
Schematic representation of dermatophytosis diagnostic methods. The methods of dermatophyte detection are summarized into two categories: (1) CRISPR-Cas12a based detection. It only takes 30 minutes after DNA extraction with high specificity and sensitivity; (2) Conventional detection methods. With Wood’s lamp, only 50% *M. canis* can be detected. Meanwhile, other foreign matters will also fluoresce, which may cause interference. Microscopic identification is a rapid diagnostic method, but it requires qualified personnel and may lead to misdiagnosis. Fungal culture is an accurate method of detecting fungal infections, but it can be time-consuming (2 weeks) and requires qualified personnel. Dermatophyte PCR and qPCR as specific molecular tools have the disadvantages of expensive equipment requirement, which brings inconvenience to clinical applications.

## 2 MATERIALS AND METHODS

### 2.1 Reagents and Instruments

The dry thermostat was purchased from Kylin-Bell Lab Instruments Co., Ltd (Haimen, China). The centrifuge was purchased from Allsheng Instruments Co., Ltd (Hangzhou, China). Thermal cycler C1000 was purchased from Bio-Rad Laboratories, Inc (America). The constant temperature incubator was purchased from LONGYUE (Shanghai, China). Fluorescence performances were measured by EnSpire Multimode Plate Reader from PerkinElmer. 2×Taq PCR Mix was purchased from TIANGEN Biotech (Beijing, China). Transcription Kit was obtained from Thermo Fisher Scientific (San Jose, CA, USA). RNAase inhibitor and NEB buffer were purchased from New England Biolabs (Ipswich, USA).

### 2.2 Clinical sample collection

We obtained six clinical samples from cats and dogs from three animal clinics in China (Table 1). The samples were collected through hair plucked out from the edges of skin lesions or with apple-green fluorescence using Wood’s lamp, and scurf gathered from hair coats. By observing different amounts of infectious materials under microscopy in the clinics, all patients were diagnosed with dermatophytosis. Through Wood’s lamp inspection, the classic apple-green fluorescence could be observed on all the hair coats of 21 patients. The patients diagnosed at China Agricultural University Veterinary Teaching Hospital (three cases in total) were not examined by Wood’s lamp (Table 1).

**Table 1.**
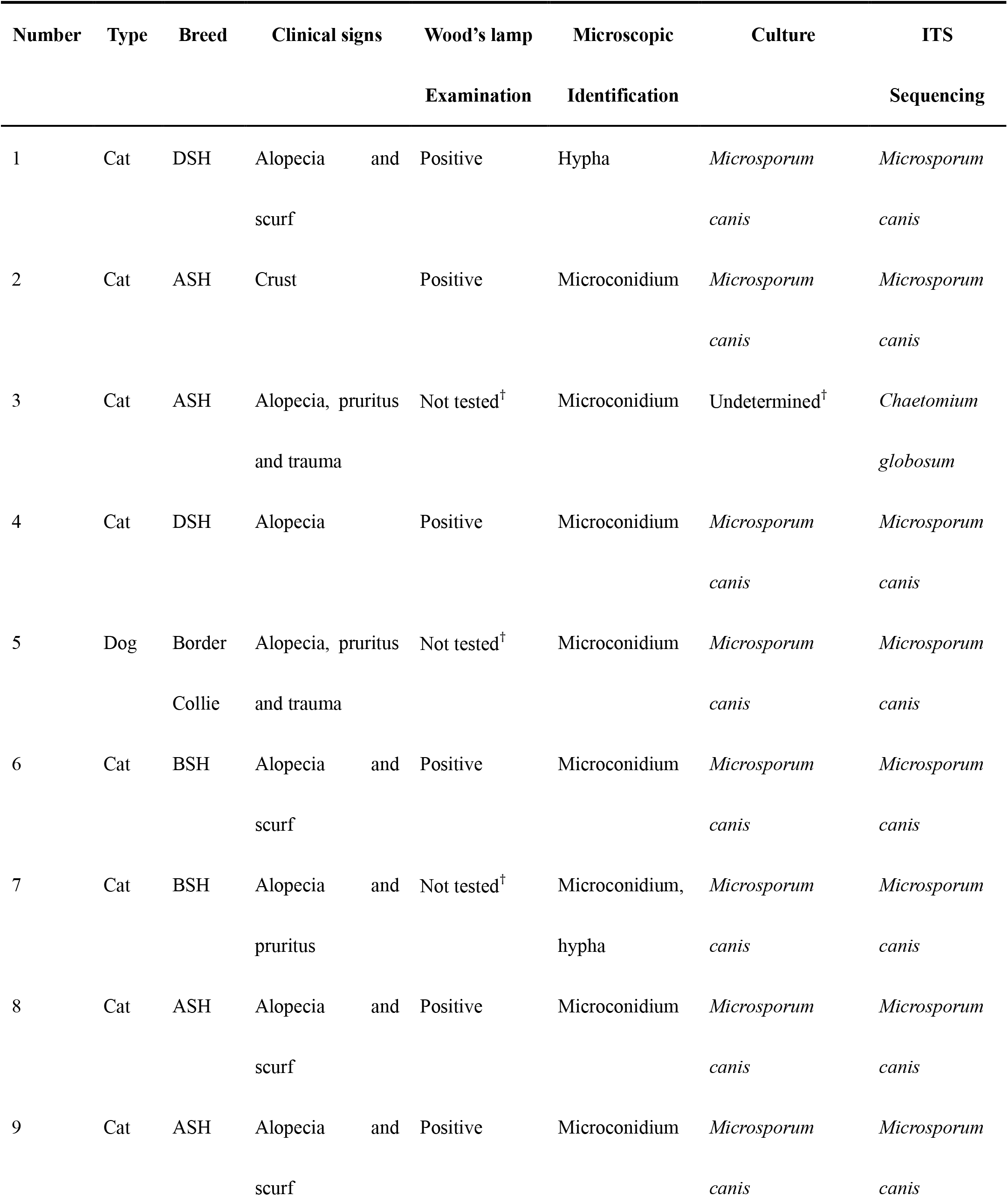
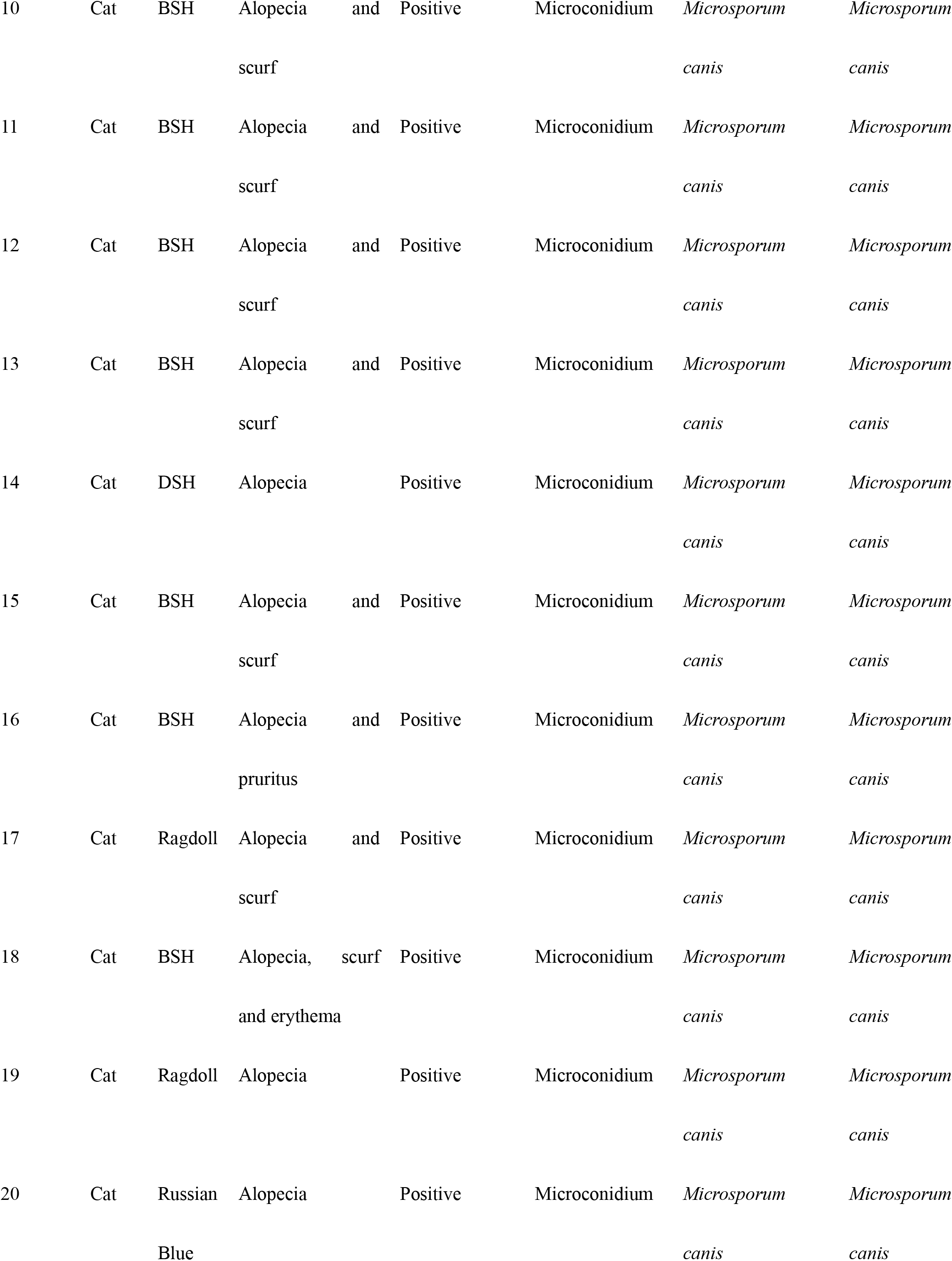
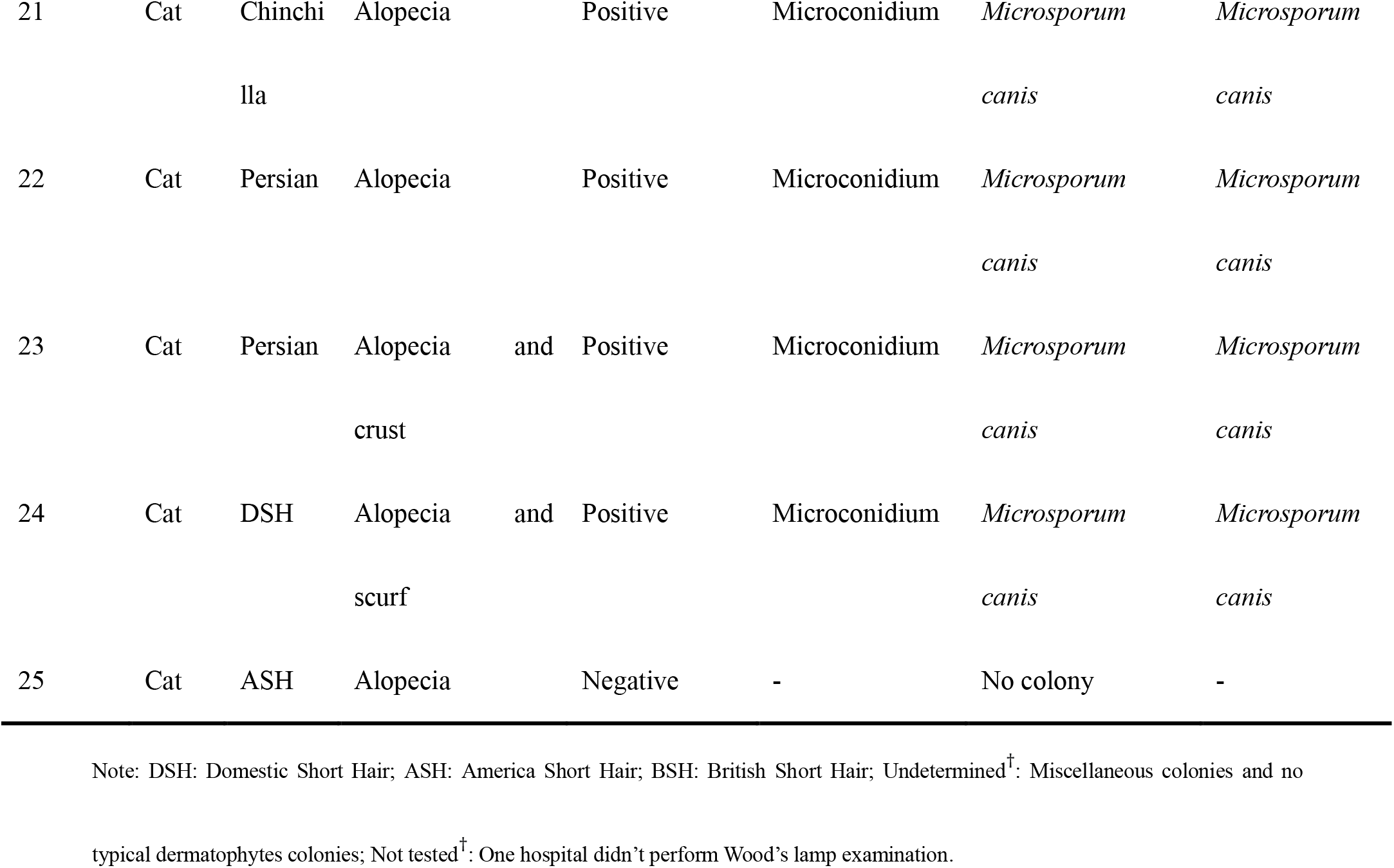
Clinical information and results of cases in the study.

### 2.3 Strains, clinical isolates and fungal culture conditions

We used the standard dermatophyte strains from American Type Culture Collection: *T. mentagrophytes* ATCC 28185, *N. gypsea* ATCC 14683 and *M. canis* ATCC 32903. In the laboratory, standard strains and clinical samples were inoculated on the Sabouraud Dextrose Agar (SDA) plate containing chloramphenicol and incubated at 30°C for 2 weeks. The samples of all cases were identified by observing the macromorphology and micromorphology of fungal colonies after about 2 weeks, and then ITS1 and ITS4 primers were used for PCR amplification. The PCR products were submitted to Sangon Biotech (Shanghai, China) for sequencing to identify species.

### 2.4 DNA Extraction from clinical samples and isolates

The hair and scurf samples of 25 animals were extracted by mixing in 45 μL extraction buffer (50 mM sodium hydroxide [NaOH]) and incubated at 95°C for 10 minutes, and neutralized by 5 μL of 1 M TRIS-HCl, pH 8.0 buffer. After mixing, the samples were centrifuged at 12,000 rpm for 5 minutes, and then the supernatant was collected. This DNA-containing solution was prepared for template for the RPA assays. The DNA in the fungal culture (pieces of colony of 3-5 mm diameter) was extracted by the same procedures as mentioned above.

### 2.5 Generation of dsDNA targets

The sequences used for PCR and RPA primers design were obtained from the NCBI Nucleotide Database (GenBank accession numbers were MH858319.1, NR_131265.1 and NR_131271.1, respectively). PCRs were performed by Q5 High-Fidelity DNA Polymerase (New England Biolabs, Inc. USA) with the listed primers (Table 2) and 2 μL genome samples, following the program: 98 °C for 30 s, then 35 cycles of 98 °C for 5 s, 58 °C for 10 s and 72 °C for 3s. Primers for RPA were listed in Table 2. RPA reactions were performed by the Twist-Amp basic kit (TwistDX, Britsh). Each RPA reaction contained 29.5 μL rehydration buffer, 2.4 μL forward and reverse primers, 2 μL genome samples, 2.5 μL of magnesium acetate (MgAc), and 11.2 μL water. The mixtures were incubated at 39°C for 15 min. Then the RPA products were cleaned up using 70% ethanol precipitation method and verified by electrophoresis on a 1% agarose gel.

**Table 2.**
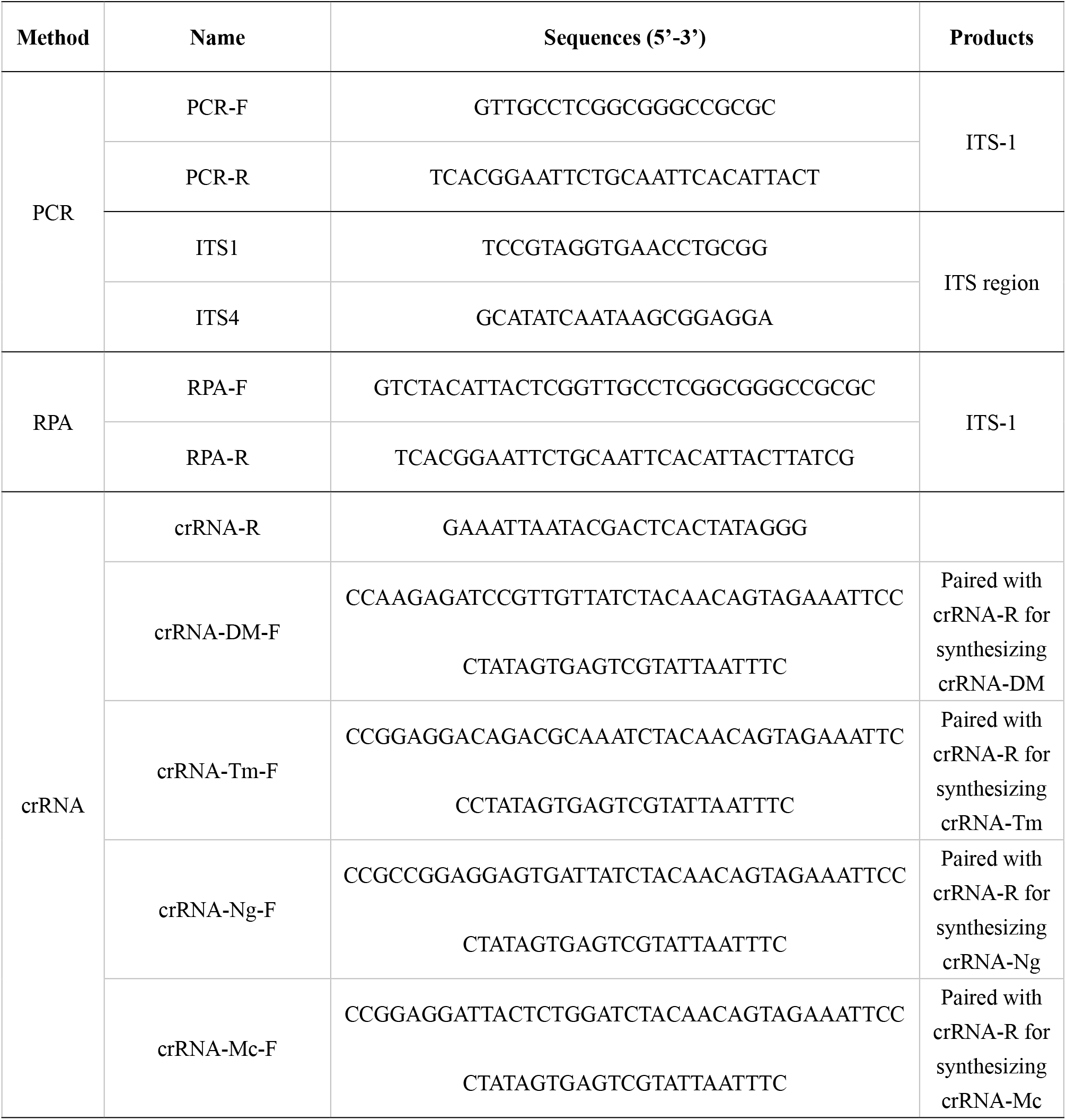
Primers used in this study.

### 2.6 Cas12a Expression and Purification

A his-tagged (C-terminal) codon-optimized version of Cas12a (*Francisella tularensis subsp. novicida*) gene was synthesized from Sangon Biotech (Shanghai, China). The expression plasmid (pET28a-FnCas12a) was transformed into BL21 (DE3), then, BL21 (DE3) cells carrying the expression plasmid were cultured in Luria-Bertani (LB) medium at 37 °C overnight. The cells were transferred into fresh LB (1:100 inoculation) at 37 °C until OD600 reached 0.8. Then, induced with 0.5 mM IPTG and transferred to 18°C for 16-hour expression. Cells were collected by centrifugation and resuspended in 50 mL of lysis buffer [50 mM Tris-HCl (pH 8.0), 1.5 M NaCl, 1 mM DTT and 5% glycerol] with 1 mM phenylmethanesulfonyl fluoride (PMSF) as the protease inhibitor and lysed by high pressure. Then the lysis was centrifuged at 15000 g for 30 min and the supernatant was loaded onto HisTrap HP column (GE Healthcare, America). The column was then washed with wash buffer (lysis buffer supplemented with 30 mM imidazole) and eluted with elution buffer (lysis buffer supplemented with 600 mM imidazole). The collected protein was dialyzed in storage buffer (20 mM Tris-HCl, pH 8.0, 600 mM NaCl, 1 mM DTT, 0.2 mM EDTA, 15% (v/v) glycerol) and finally stored in aliquots at −80 °C.

### 2.7 Transcription of crRNAs

The preparation crRNA was proceeded in three steps. The transcription templates for crRNA preparation were amplified by the PCR process, and the primers were list in Table 2. Then, the transcription process was performed at 37°C overnight using the T7 High Yield Transcription Kit (Thermo Fisher Scientific, USA). Finally, the transcript products were purified using the RNA Clean & ConcentratorTM-5 (Zymo Research, USA and quantified with NanoDrop 2000C (Thermo Fisher Scientific, USA).

### 2.8 Cas12a detection

Cas12a cleavage reaction system contained 500 nM Cas12a, 500 nM crRNA, 2 μL target DNA, 500 nM ssDNA (FAM-GATCAAGAGCTA-BHQ1) and 0.5 μL RNase inhibitor (TaKaRa, Japan) in a 20 μL volume. The reactions were performed at 37°C in NEB buffer 3.1 for 15 min, and the reaction products in a volume of 10 μL were diluted 10 times to 100 μL and examined using EnSpire Multimode Plate Reader (PerkinElmer, America). Then, the remaining reaction products were placed under the blue light generated by Azure C300 Gel Imager (Azure Biosystems, America) for visual inspection.

## 3 RESULTS

### 3.1 Design and detection of crRNA guides and primers in CRISPR-Cas12a assay

Internal transcribed spacer (ITS) region was selected for RPA-Cas12a detection, because of their divergent sequences across different fungi species and high probability of identifying the fungi type. Based on the sequence alignment of ITS1 regions, four crRNA guides were designed to identify the three major dermatophytes infections in cats and dogs. A specific sequence was selected for crRNA guide recognition of three main <u>d</u>er<u>m</u>atophytes (crRNA-DM) and three specific sequences were selected for crRNA guide recognition of three specific dermatophytes (crRNA-Ng, Tm, and Mc) (Fig. 2A). PCR amplification was performed with DNA extracted from the standard strains by using primer set (PCR-F and PCR-R in Table 2) from fungal ITS1 region. The target products were amplified from extracted DNA from all three strains (Fig. 2B).

**Fig. 2.**
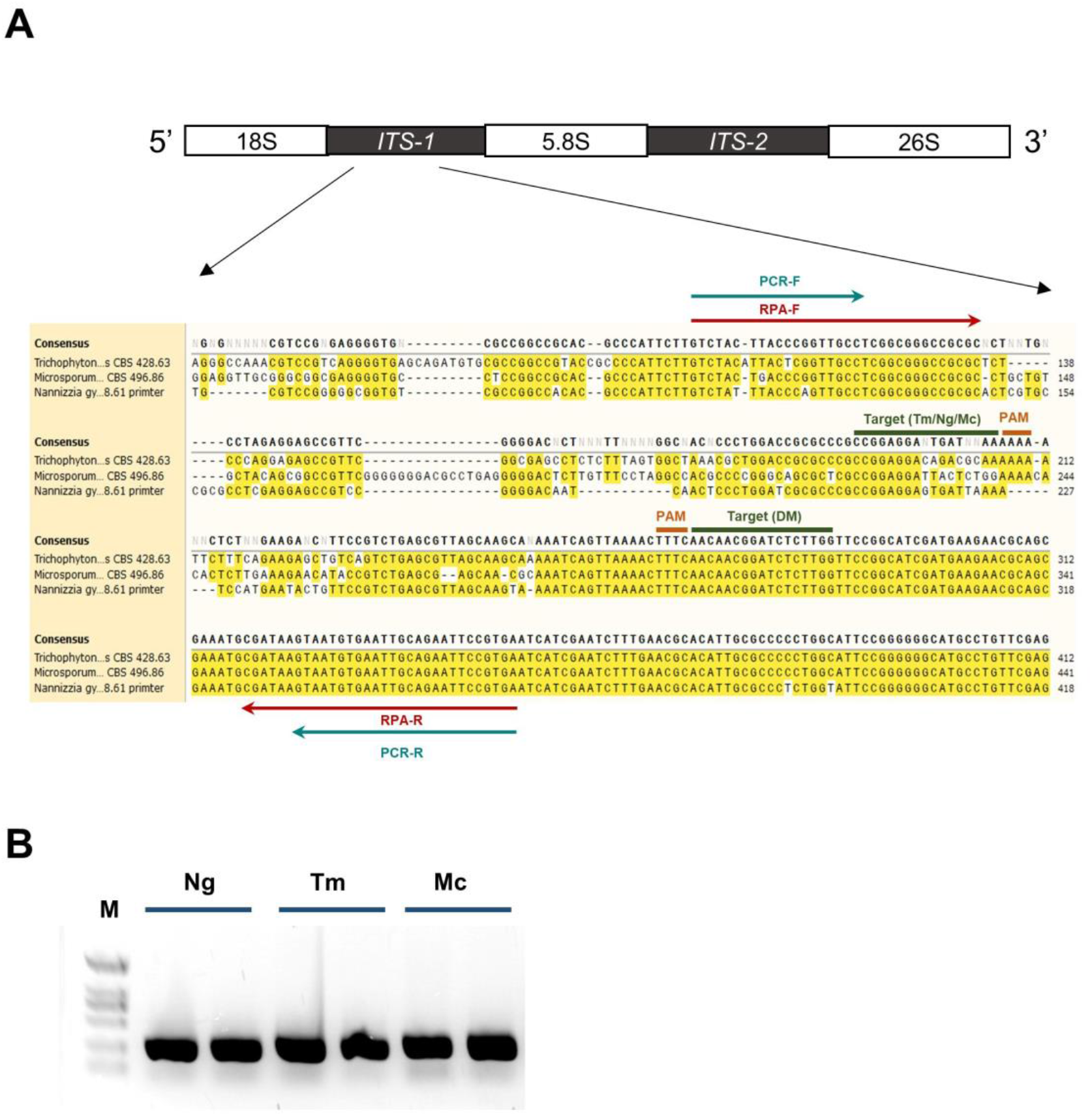
Primers and crRNA locations on ITS1 region. **A**, Green lines indicate the primers of PCR; red lines indicate the primers of RPA; target sites of the crRNAs were marked in the corresponding area. The sequence alignment was performed by SnapGene software. **B,** *N. gypsea*, *T. mentagrophytes* and *M. canis* standard strains PCR products analysis. M: molecular size marker (Target fragment size 210 bp); results of PCR performed for *N. gypsea* (lane 1 and lane 2), *T. mentagrophytes* (lane 3 and lane 4) and *M. canis* (lane 5 and lane 6).

4 (Fig. 3A). Under visual inspection of the reacted solutions, fluorescence signals of all three standard samples can be observed. Then, we applied three specific crRNAs for identification of corresponding dermatophytes. The fluorescence detection using crRNA-Ng guide can specifically identify *N. gypsea*, and compared with the negative control, it has reached a significant difference (p<0.001). However, fluorescence detections of the remaining dsDNA products (*T mentagrophytes* and *M. canis*) were higher than the negative control, but they can’t reach a significant difference (p<0.001) (Fig. 3B). Therefore, we set the “p<0.001” as the detection standard and only significant values (p<0.001) were marked with asterisks (***). The guides crRNA-Tm and crRNA-Mc also can specifically identify *T. mentagrophytes* and *M. canis*, respectively with significant differences (p<0.001) (Figs 3C and 3D). Under visual inspection, the target can produce obvious and specific fluorescence in blue light (Figs 3E–3H).

**Fig. 3.**
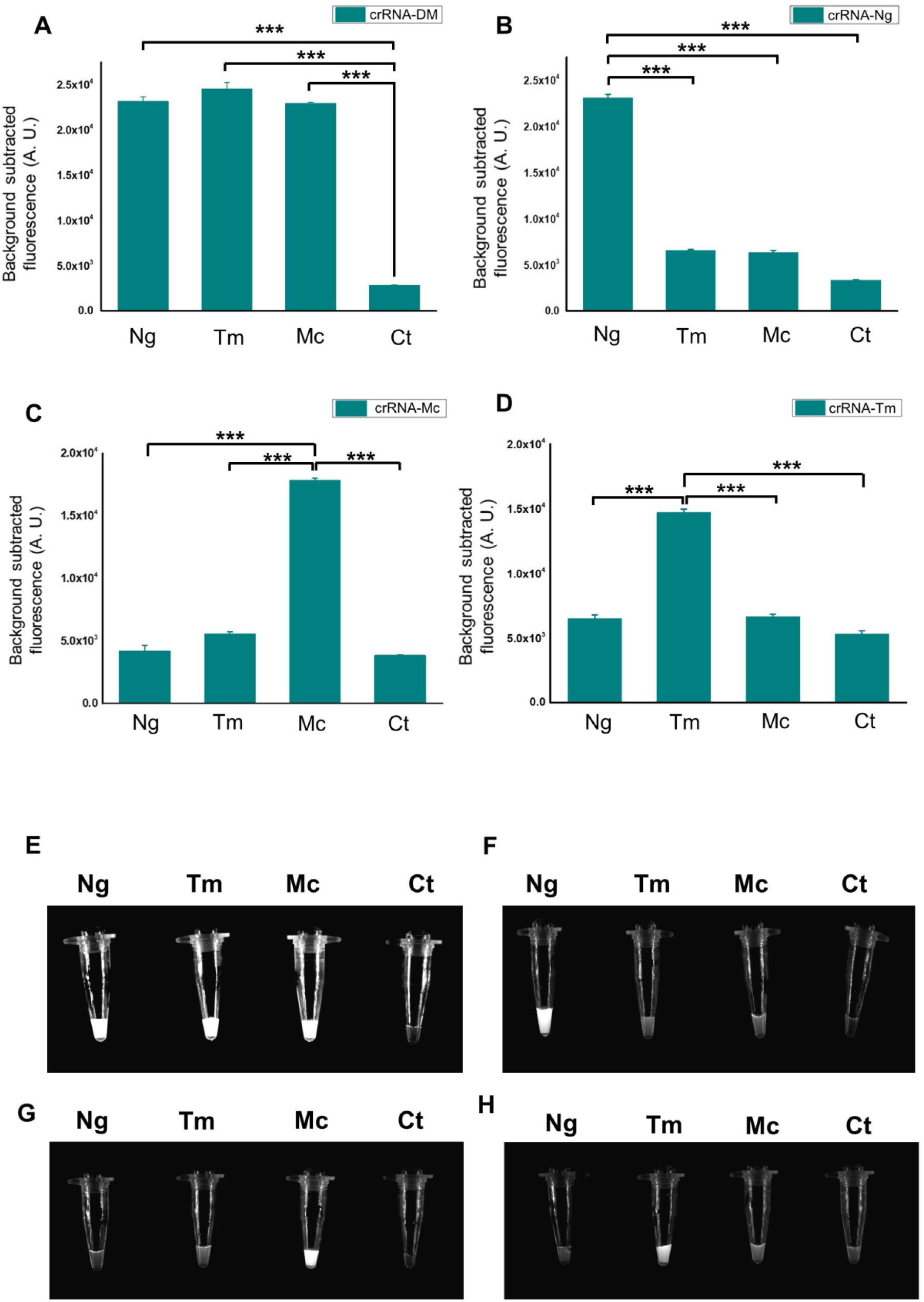
Four specific guides designed for CRISPR-Cas12a assay. **A-D,** Fluorescence detection of *N. gypsea* (Ng), *T. mentagrophytes* (Tm) and *M. canis* (Mc) using guides crRNA-DM, crRNA-Ng, crRNA-Tm and crRNA-Mc respectively. **E-H,** Visual detection of *N. gypsea* (Ng), *T mentagrophytes* (Tm) and *M. canis* (Mc) using guides crRNA-DM, crRNA-Ng, crRNA-Tm and crRNA-Mc under blue light. n = 3 technical replicates, bars represent mean±SD, statistical analysis was performed using one-way ANOVA test and only significant (p < 0.001) values were marked with asterisks (***): ***p <0.001. CT, negative control which is without the target DNA.

### 3.2 Sensitivity of the Cas12a-based fluorescence reporting system

In order to determine the limit of detection (LOD), different concentrations of dsDNA products were participated in the Cas12a-based detection. Signals generated by the use of guides crRNA-Tm and crRNA-Mc showed significant differences (p<0.001) between experiment groups with 10 nM DNA concentration and the negative control (Figs 4A and 4B). Compared to the control, there were no significant difference when the concentration of the samples was less than 10 nM (Figs 4A and 4B). In parallel, signals generated by the use of guides crRNA-DM and crRNA-Ng showed significant values (p<0.001) by using 100 nM DNA concentration in statistical analysis, and there were no significant differences when the concentration of the samples were less than 100 nM (Figs 4C and 4D). The results indicated that Cas12a assay can detect nucleic acid products with a concentration as low as 10-100 nM.

**Fig. 4.**
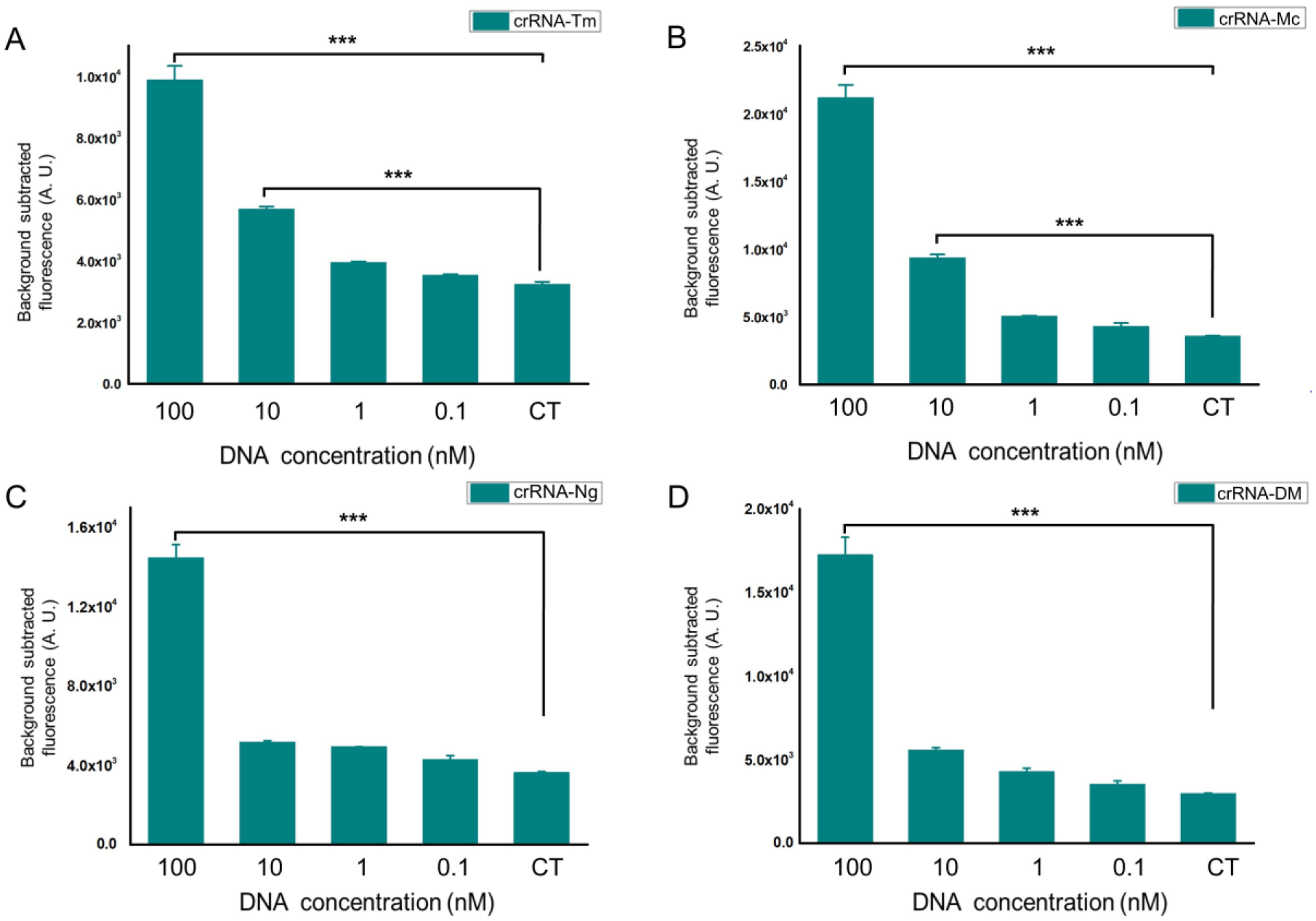
Sensitivity of the Cas12a-based fluorescence reporting system. **A,** Fluorescence detection using guide crRNA-Tm. **B,** guide crRNA-Mc. **C,** guide crRNA-Ng. **D,** guide crRNA-DM. Cas12a assay can detect as low as 10 nM of PCR products. n = 3 technical replicates, bars represent mean± SD, ***p <0.001. CT, negative control which is without the target DNA.

Combined with the RPA reaction, a copy number gradient of per RPA reaction (10^0^, 10^1^, 10^2^, 10^3^, 10^4^, 10^5^, 10^6^ and 10^7^) was tested for sensitivity examination. We found that whatever copy number was used, all detection assays showed significant values (p<0.001) between experimental group and negative control in statistical analysis, which were marked with asterisks (***). Under visual inspection, the results were the same. The significant fluorescence changes can be observed and were different from the negative control (Fig. 5). The findings suggested that using RPA coupled with Cas12a-based assay can detect as low as a single copy of genome.

**Fig. 5.**
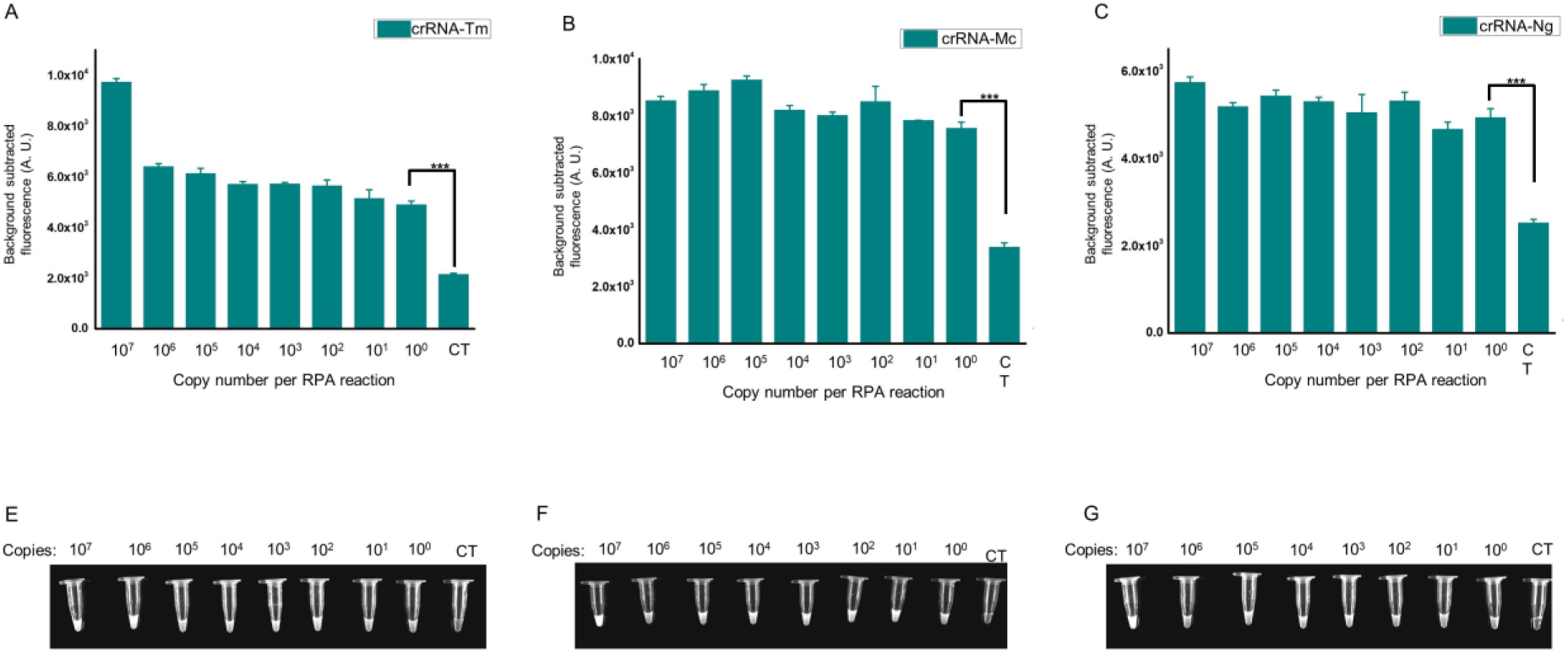
Detection of dermatophytes using RPA coupled with Cas12a-based assay. **A-C**, Fluorescence detection of RPA products. Reactions can detect as few as a single copy of genome. **D-E**, Direct observation by naked-eyes when the Cas12a reactions system exposed to blue light. n = 3 technical replicates, bars represent mean± SD, ***p <0.001. CT, negative control which is without the target DNA.

### 3.3 The specificity and sensitivity of RPA-Cas12a fluorescence assay for clinical samples

25 clinical samples were collected for RPA-Cas12a assay verification. The clinical diagnosis of dermatophytosis was based on clinical signs, Wood’s lamp examination and direct microscopy. All samples had different degrees of clinicals appearances, and infectious materials can be found under the microscope (Table 1), of which 24 cases were diagnosed with dermatophytosis by veterinarians. The case No. 25 diagnosed as non-dermatophytosis was collected as negative control. This patient showed focal hair loss which was similar to the signs of dermatophytosis and the clinical diagnosis result was non-dermatophytosis. We used the RPA-Cas12a fluorescence assay to detect clinical samples, so that we can verify the specificity and sensitivity of this new rapid diagnostic method. The fungal culture and sequencing were considered as standard results.

To evaluate our method, we initially tested our method using the first 6 clinical samples. The crRNA-DM guide can detect fluorescence of 5 samples and identify them as dermatophytes infections, which reached significant differences (p<0.001) except for sample No. 3 (Figs 6A and 6B). The crRNA-Mc guide also detected obvious fluorescence in these 5 samples (Figs 6A and 6B). In addition, crRNA-Ng and crRNA-Tm guides can not detect any fluorescence from these 6 samples. Then we collected and tested more samples. Sample No. 7 to No. 24 were all identified as *M. canis* (Figs S1 and S2A). We were surprised about the results for sample No. 3, as it was initially diagnosed with dermatophytosis in clinics. We inoculated all 25 clinical samples on SDA plate and incubated at 30°C for 2 weeks, and all samples were grown expect for negative control sample No. 25 (Figs 6C and S2B). Then we performed micromorphology study and ITS sequencing using ITS1 and ITS4 as primer set for these clinic isolates. After observing the macromorphology of fungal colony, micromorphology of macroconidia and performing NCBI Blast of ITS sequencing, 23 samples (except for samples No. 3) were identified as *M. canis* (Figs 6C and S2B). The RPA-Cas12a fluorescence test matched the fungal culture and ITS sequencing results. The result of negative control (No. 25 sample) was also consistent with the results of fungal culture and clinical diagnosis (Fig. S2B). After statistical analysis, RPA-Cas12a method can achieve 100% sensitivity and 100% specificity (Table 3). The main colony of sample No. 3 in the medium was sequenced and identified as *Chaetomium globosum*, which is a saprophytic fungus (not dermatophytes).

**Fig. 6.**
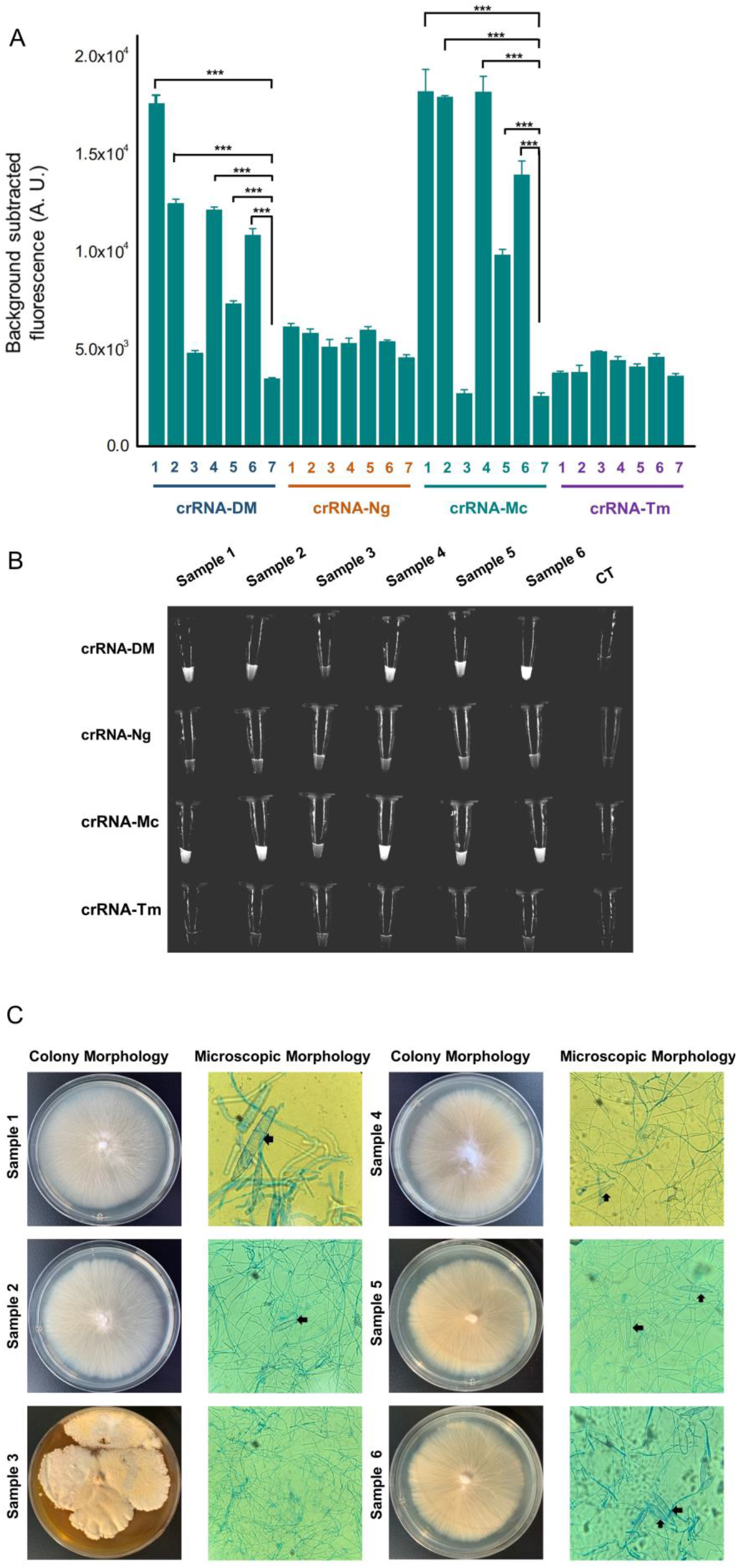
Clinical cases diagnosis by RPA-Cas12a. Collect animal hair or scurf to extract nucleic acid, and then use the nucleic acid as a template to amplify target genes by RPA. Finally, observe the fluorescence signals excited by the Cas12a reactions or detect the fluorescence signals by naked-eyes under blue light. **A**, Fluorescence detection of six samples using guide crRNA-DM, crRNA-Ng, crRNA-Tm and crRNA-Mc respectively. n = 3 technical replicates, bars represent mean ± SD, statistical analysis was performed using one-way ANOVA test and only significant (p < 0.001) values were marked with asterisks (***): ***p <0.001. **B**, Visual detection by naked-eyes under blue light. CT, negative control which is without the target DNA. **C**, Fungal colonies of 5 samples were flat, spreading, cream-colored, with a dense cottony surface and bright yellow reverse pigment. The appearances were consistent with *M. canis*. Under the microscope, these samples had the spindle-shaped macroconidia (arrow: macroconidia) with 5-15 cells and a terminal knob, consistent with *M. canis*. The sequencing results proved that pathogen of these samples was *M. canis*. Both the colony and microscopic morphology of sample No. 1-3 were inconsistent with the characteristics of dermatophytes, 400× magnification.

**Table 3.** Performance of Cas12a assay for diagnosis of dermatophyte.

|  | Culture positive | Culture negative | Total |
| --- | --- | --- | --- |
| Cas12a positive | 23 | 0 | 23 |
| Cas12a negative | 0 | 2 | 2 |
| Total | 23 | 2 | 25 |
| Sensitivity |  | 100 |  |
| Specificity |  | 100 |  |

Note: Values are shown as %

## 4 DISCUSSION

At present, the identification of dermatophytes in small animals still relies on comprehensive diagnosis methods, such as Wood’s lamp examination and microscopy in clinical practice. Some clinics can perform fungal culture and dermatophyte PCR for further diagnosis. However, not every clinic has relative equipment that can be used and the fungal culture will delay the diagnosis for about 1-2 weeks. Actually, there is no gold standard diagnostic test for dermatophytosis.^3^ Every diagnosis method used clinically has its own limitations. For example, hair loss, scaling, crusting and erythema are the most common clinical lesions. Due to the variability of the lesions, pruritus and other lesions may occur in different animals. Moreover, it also can be confused with other diseases such as pemphigus foliaceus. As for Wood’s lamp examination, only 50% of *M. canis* infections can be detected with fluorescence, while other dermatophytes (such as *N. gypsea* or *M. persicolor*^32^) do not produce fluorescence.^33^ It can cause confusion and inaccuracy at the same time.^2^ Direct microscopic examination may be a simple and rapid way to screen for microconidia and hyphae, however, only >85% of cases can be accurately diagnosed.^3^ In addition, the professional requirements for microscopic examination are very high. Two recognized disadvantages of fungal culture are time-consuming and demanding (about 1-2 weeks), which can delay the diagnostic outcome and treatment. Molecular tools are increasingly used in the laboratory for the fungal identification. Dermatophyte PCR or qPCR is becoming more and more popular due to its sensitivity, but it is not intuitive and more dependent on available laboratory conditions and the facilities, and the processing time is about 2-3 hours.

Accurate and timely identification of fungal isolates are very important. All of these emphasizes the actual need to develop new methods that provide rapid, intuitive and highly specific identification of dermatophytes. We developed an RPA-Cas12a detection method for dermatophytes, which showed excellent results in our research. The results were 100% consistent with the traditional fungal culture and ITS sequencing. We found that no matter what kinds of samples we use to extract DNA, such as hair, scurf or the debris on the tape or cotton swab, the amount of extracted DNA was sufficient for later Cas12a-based diagnosis. The Cas12a-based assay can detect as low as 10-100 nM PCR product, which showed that the developed assay had high sensitivity. All the positive samples can be sorted and visualized with naked-eyes within 30 minutes at a constant temperature, which took less time than other molecular methods. Meanwhile, it was easy to implement in both basic laboratories and on-site detection.

In our study, one sample (sample No. 3) could only be screened out a few spores under the microscope in clinics and then could not be identified by CRISPR-Cas12a assay. In the fungal culture, no typical dermatophyte colonies grew. The main colony in the medium was ITS sequenced and identified as *Chaetomium globosum*, which is a saprophytic fungus. It primarily resides on plants, soil, straw and dung, which has not been shown as pathogen. Based on the negative result of the culture, we suspected that the saprophytic fungi spores observed under the microscope may be carried by the animal patient, and this caused a misdiagnosis. The clinical signs may be caused by other reasons in the patient while *Chaetomium globosum* could be the result of environmental contamination. Therefore, the method we designed can assist veterinarians in accurately diagnosing three main dermatophytosis, especially when the operators are not yet proficient in the microscopic examination. In our study, we only used 25 clinical cases for verification. Because of regional differences and different pet-raising styles, there are relatively few cases of dermatophytosis in small animals caused by *N. gypsea* and *T. mentagrophytes*, especially in cats^3^. Therefore, more specimens should be included in the experiment to evaluate the effectiveness in the future.

In conclusion, the RPA-Cas12a-fluorescence assay is a promising method for detecting dermatophytes with high sensitivity and specificity. And this method can rapidly detect the dermatophyte genome from clinical samples onsite, and adopt optimized steps to reduce the detection time. Therefore, the RPA-Cas12a-fluorescence assay will be an excellent choice for point-of-care dermatophytosis diagnosis.

## ACKNOWLEDGEMENTS

We thank Degui Lin’s lab (Clinical Department, College of Veterinary Medicine, China Agricultural University) for providing the standard strains, and China Agricultural University Veterinary Teaching Hospital, Guangzhou United Animal Hospital, Chang An Pet Hospital, Stitch Veterinary Hospital and Guangzhou Tayan Animal Hospital for providing the clinical samples of fungi. We are also grateful to the pet owners for their cooperation in sampling.

## AUTHOR CONTRIBUTIONS

**Liyang Wang:** data curation (equal); investigation (equal); methodology (equal); writing-original draft (equal); formal analysis (equal). **Jinyu Fu:** data curation (equal); investigation (equal); methodology (equal); writing-original draft (equal); formal analysis (equal). **Guang Cai:** investigation (equal); methodology (equal). **Di Zhang:** writing-review and editing (equal); investigation (equal). **Shuobo Shi:** writing-review and editing (equal); supervision (equal); validation (equal). **Yueping Zhang:** writing-review and editing (equal); supervision (equal); validation (equal).

