## Supplementary for "Rapid and Visible RPA-Cas12a fluorescence Assay for Accurate Detection of Zoonotic Dermatophytes"

### Supplementary Figures

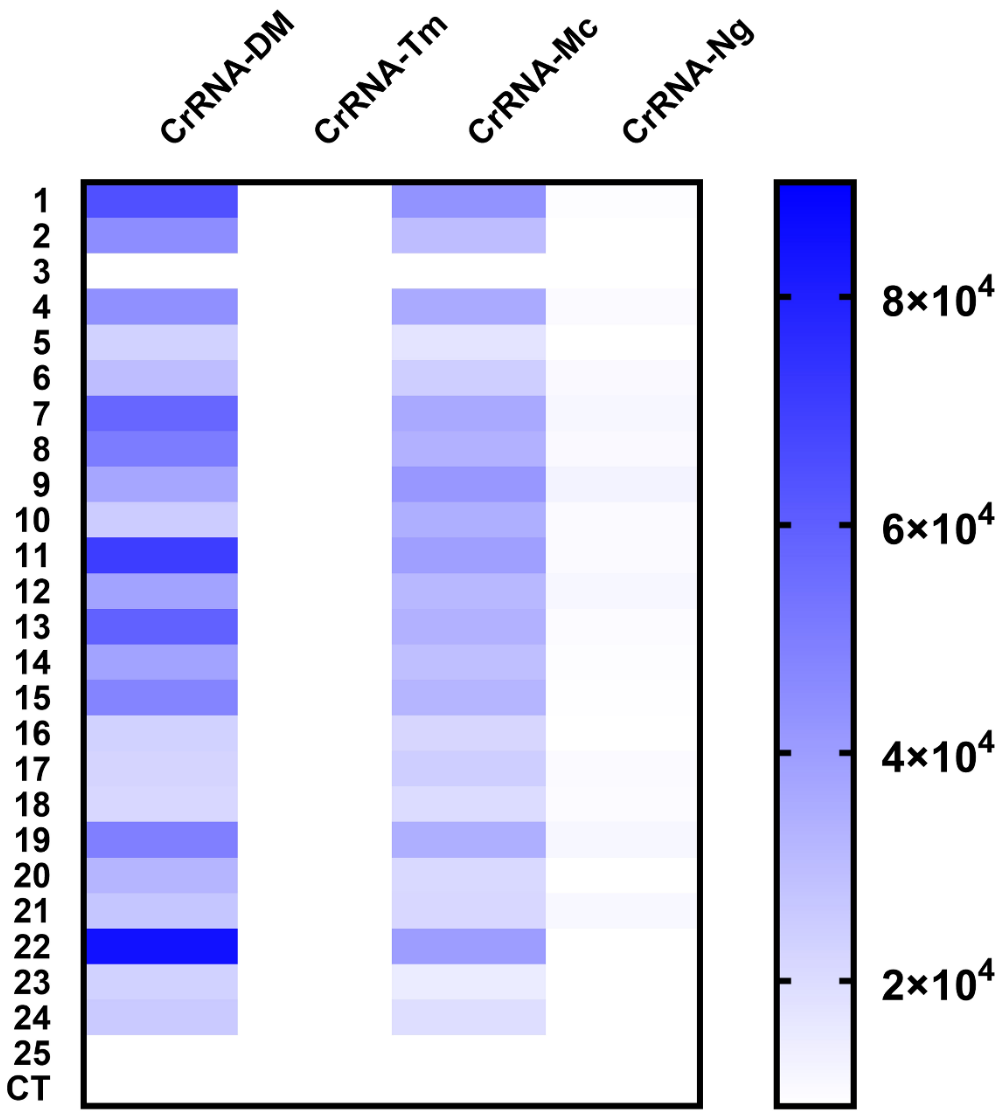


**Fig. S1 Heatmap of clinical samples by RPA-Cas12a.** Fluorescence detection of 25 samples using guide CrRNA-DM, CrRNA-Ng, CrRNA-Tm and CrRNA-Mc respectively. 23 samples were identified as *M. canis*. CT, negative control which is without the target DNA. Sample No. 25 was negative control collected from clinics.

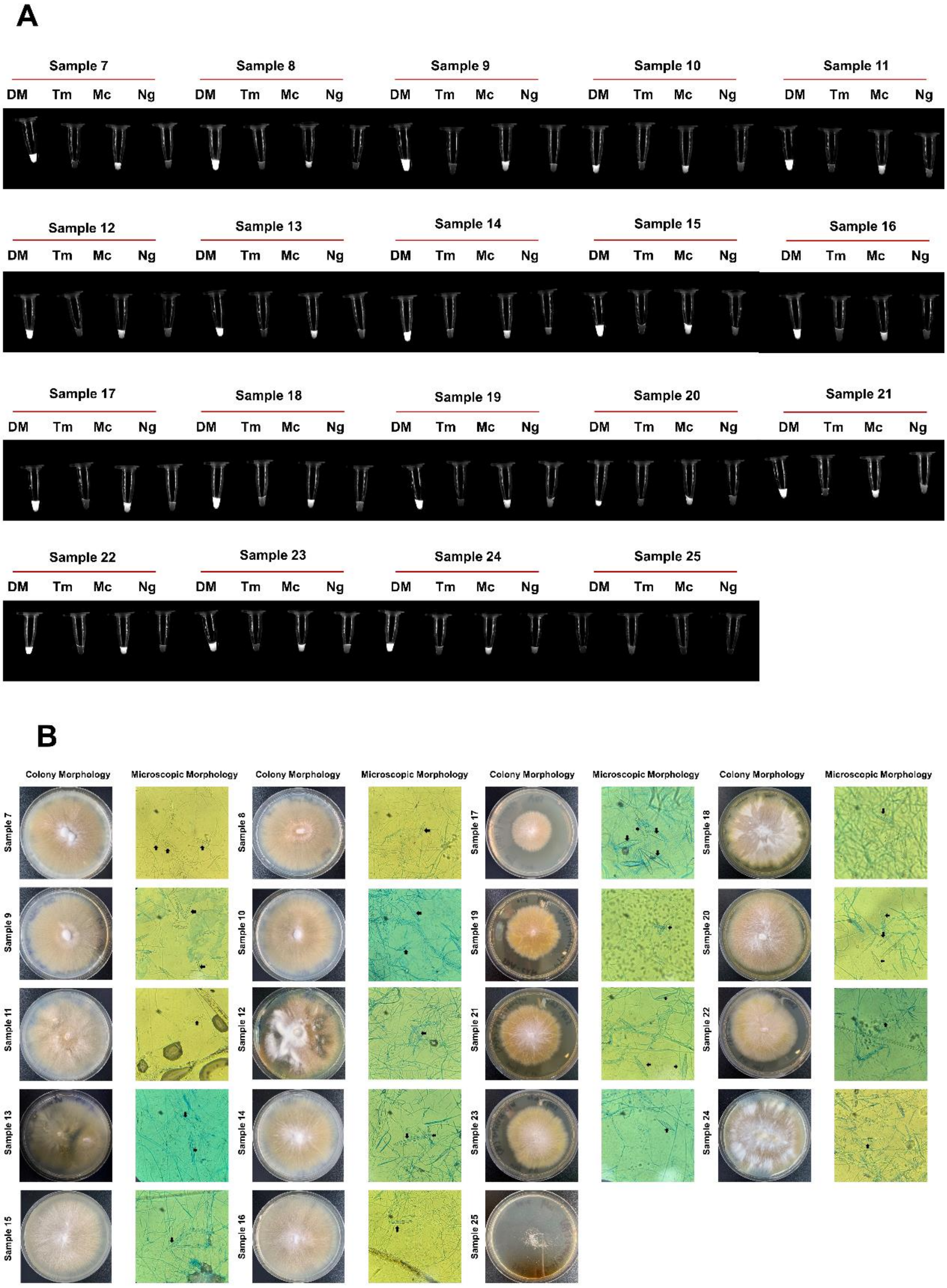


**Fig. S2 Expanded clinical samples diagnosis by RPA-Cas12a. A**, Visual detection by naked-eyes under blue light. n = 3 technical replicates. CT, negative control which is without the target DNA. **B**, Fungal colonies of 23 samples were flat, spreading, cream-colored, with a dense cottony surface and bright yellow reverse pigment. The appearances were consistent with *M. canis*. Under the microscope, these samples had the spindle-shaped macroconidia (arrow: macroconidia) with 5-15 cells and a terminal knob, consistent with *M. canis*. The sequencing results proved that pathogen of these samples was *M. canis*, 400× magnification. Sample No. 25 was negative control collected from clinics.
